# *E. coli* associated with Crohn’s disease exhibit distinct strategies to colonize macrophages

**DOI:** 10.1101/2024.09.04.611154

**Authors:** Emma Bruder, Hosni Nedjar, Nicole Quenech’Du, Caroline Chevarin, Emilie Vazeille, Marie Granotier, Parul Singh, Anthony Buisson, Nicolas Barnich, Olivier Espéli

**Affiliations:** CIRB – Collège de France, CNRS, INSERM, Université PSL, Paris, France; Microbes, Intestin, Inflammation et Susceptibilité de l’Hôte. UMR Inserm/ Université Clermont -Auvergne U1071, USC INRAE 1382, Clermont-Ferrand, France

## Abstract

Patients with Crohn’s disease exhibit abnormal colonization of the intestine by Proteobacteria, particularly the adherent-invasive *Escherichia coli* (AIEC) group. These bacteria are predominant in the mucus, adhere to epithelial cells, colonize them, and survive inside macrophages. We recently demonstrated that the acclimation of strain LF82 to phagolysosomal stress occurs in two distinct steps: first, a replication halt producing stress-tolerant persisters, and second, a replication phase that leads to the formation of Intracellular Bacterial Communities (IBC) organized with a biofilm-like matrix. Given the significant genomic diversity among strains with the AIEC phenotype, we conducted a comparative analysis of the genomes and macrophage colonization characteristics of 13 AIEC strains collected from patients during a clinical study conducts by the CHU of Clermont-Ferrand. Our results demonstrate that IBCs serve as replicative niches for all AIEC strains within macrophages. However, these strains form IBCs using different strategies, including varying levels of phagosome detoxification, distinct biofilm characteristics, and diverse macrophage responses. Our study reveals a strong positive correlation between vacuole acidification and persister induction that explains intracellular survival of the different strains. In addition, we revealed distinct AIEC dissemination strategies outside macrophages, which may contribute to the propagation of inflammation in the human host. These findings highlight that research on pathogens and pathobionts with plastic genomes should not rely solely on a few laboratory models.

## Introduction

*E. coli* strains isolated from patients suffering from Inflammatory Bowel Disease (IBD) are frequently tested for a potential AIEC phenotype, i.e. adherence and invasion to epithelial cells and survival for 24h within macrophages. AIEC prevalence is particularly high in Crohn disease (CD) and Ulcerative Colitis (UC) (López-Siles M et al., 2022). These tests are standardized and the attribution of the AIEC phenotype is based on quantitative indicators (Darfeuille-Michaud A et al., 2004). As other intestinal pathogenic *E. coli* are extracellular pathogens, the intracellular colonization is a particularly intriguing trait of AIEC (Mansour S et al., 2023; Zangara MT et al., 2023). Recent animal model experiments demonstrated that AIEC phenotypes not only play an ecological role in promoting inflammation but also contribute to disease pathogenesis (Sheikh IA et al., 2024). Strong associations between *E. coli* survival and replication in macrophages, epithelial cells *in vitro*, and strain pathogenicity *in vivo* were observed (Kittana H et al., 2023). Moreover, it has now been demonstrated that, compared to healthy individuals, macrophages in the intestines and *lamina propria* of CD patients host many more bacteria. Among them a large number of Proteobacteria were found in this dysbiotic flora (Dheer R et al., 2020). Such immune tolerance may contribute to the disease by sustaining inflammation. Hence, the necessity to better understand AIEC as a putative instigator or propagator of the disease is certain.

Based on LF82 observations, the current perception of AIEC intracellular lifestyle is that they reside in mature phagolysosomes and endure stresses brought by lysosomes such as acidic pH, oxidative molecules, toxic compounds (metals or antimicrobial peptides) and nutrient depletion (Zangara MT et al., 2023). Their eventual dissemination outside of this environment has not yet been described. We demonstrated that for LF82, the reference strain of the AIEC group (Boudeau J et al., 1999; Glasser AL et al., 2001), the colonization of macrophage results from a sophisticated strategy involving core genome responses, activation of horizontally acquired pathogenicity islands, phenotype switches and the formation intracellular communities within vacuoles (Demarre G et al., 2019; Prudent V et al., 2021). Interestingly, within macrophages LF82 becomes antibiotic tolerant (Bruder E & Espéli O, 2022; Demarre G et al., 2019). The strategy used by LF82 differs drastically from textbook’s description of macrophage colonization by classical models such as *Salmonella*, *Brucella*, *Legionella* or *Mycobacteria* that will eventually counteract immune response by secreting virulence factors.

AIEC identification is currently challenging because it relies on phenotypic assays based on infected cell cultures. To address this issue, a search for AIEC molecular markers has started; however, a specific and widely distributed genetic AIEC marker is still missing (Camprubí-Font C & Martinez-Medina M, 2020). Although different groups have analyzed the genome content of AIEC strains (Camprubí-Font C et al., 2018; Rakitina DV et al., 2017), the complete and annotated genomes of only three AIEC strains (LF82, NRG857c and UM146) are currently available in the NCBI database (Krause DO et al., 2011; Miquel S et al., 2010; Nash JH et al., 2010).

Because intestinal colonization, adherence and invasion of epithelial cells are properties shared by *E. coli* pathogens, we postulated that intra-macrophage colonization might be a genuine marker of AIEC. Therefore, in the present study, we combined genomic analysis of several AIEC strains with in depth description of their macrophage colonization strategies. We generated draft genomes from 10 new AIEC strains. We assembled and annotated the complete genome of 4 new AIEC strains belonging to three different *E. coli* phylogroups. We demonstrated that all AIEC form intravacuolar communities (IBC) that are characteristic of AIEC compared to other enterobacterial pathogens and commensals. Our study also showed that persistent IBC can be formed using different pathways and may lead to different macrophage fates. Altogether, these data illustrate that for AIECs, the study of one or few reference strains is not enough to understand host colonization strategies and subsequently fight infections. Probably, for all the bacterial strains that possess significant genomic plasticity, such as ESKAPE, the study of several strains seems to be necessary to understand the mechanisms of infection.

## Results

### Phylogenetically distant AIEC strains colonize macrophages

To delineate strategies for macrophage colonization, we gathered *E. coli* strains exhibiting the AIEC phenotype (adhesion, invasion of epithelial cells and survival within macrophages, Fig. 1A and Supplementary Fig. S1A). These strains were collected from CD patients biopsies in the course of two different clinical multicentre studies (Buisson A et al., 2021; Martinez-Medina M et al., 2009). While draft genomes were previously available for some strains (Camprubí-Font C et al., 2018), we conducted sequencing for the remaining strains. Annotation of the assembled contigs unveiled significant genomic diversity within this group (Fig. 1B and 1C). Core genes represent 43% of the pangenome (7696 gene clusters), soft core genes account for 10%, shell genes for 30%, and singleton genes for 17%. Notably, certain strains harbor over 200 singletons gene clusters that are absent in all other strains. Consistent with this diversity, AIECs from our collection were assigned to four of the six *E. coli* phylogroups (A, B1, B2, and D). We assessed the strains’ capacity to colonize macrophages by quantifying colony-forming units (CFU) at 24 hours post-infection (Fig. 1A) and evaluating their ability to form intra- vacuolar communities (Fig. 1D and Supplementary Fig. S1B). For the majority of strains tested, macrophage survival is robust but the colonization is not extensive (with a 24-hour to 1-hour ratio comprised between 0.5 and 2). Notably, strains from the B2 group exhibit higher survival rates (median survival rate = 1.14) compared to those from the A (median survival rate = 0.62; p-value = 0.012) and D (median survival rate = 0.46; p-value = 0.003) groups. Moreover, survival rates were significantly greater than those of the IAI4 commensal strain, the UPEC CFT073 strain and *Shigella flexneri*, yet lower than survival rate of *Salmonella Tiphymurium* (Fig. 1A). Communities of bacteria residing within Lamp-I positive vacuoles were consistently observed for each AIEC strain (as depicted in Fig. 1D and Supplementary Fig. S1B). These communities comprised tens to hundreds of bacteria densely packed within a single vacuole (Supplementary Fig. S1C). It’s noteworthy that a single macrophage could harbor multiple vacuoles teeming with AIECs. Interestingly, these AIEC communities exhibit stark differences from those associated with IAI4, which are notably smaller and less frequent. Furthermore, they differ from those formed by CFT073, *Salmonella*, or *Shigella*, which display smaller carrying capacities (typically less than 10 bacteria per vacuole, Fig. 1D). These observations underscore the significance of intra-vacuolar community formation as a defining characteristic of AIECs, despite their diverse genomic contents.

**Figure 1:**
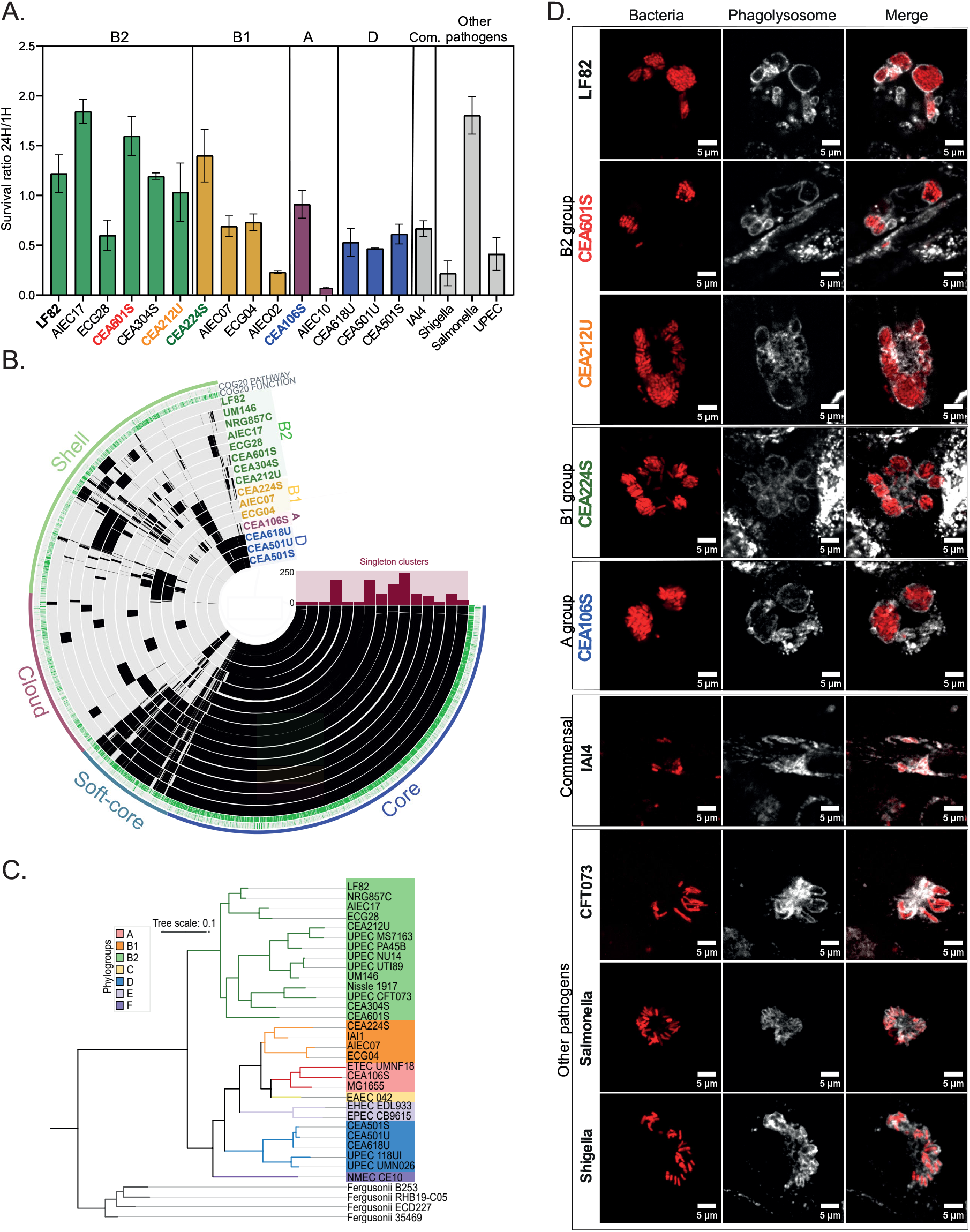
In spite of different genome contents every AIEC colonize macrophage by forming IBC. A) Phylogenetic tree of AIEC strains, other pathogenic *E. coli*, and commensal strains, based on pangenome genes. The tree was unrooted using *E. fergusonii* strains. B) Pangenome analysis of 14 AIEC strains generated with Anvi’o. The layers represent individual genomes organized by their phylogenomic relationships. In the layers, black indicates the presence of a gene cluster, while light colors indicate its absence. The colors of the labeled genes correspond to different phylogroups within *E. coli*. C) "Survival of the 13 AIEC strains, 2 strains (ECG04 and ECG28) closely related to AIEC (differing by only 3 SNPs), IAI4 (a commensal strain), and 3 pathogenic strains (Shigella, Salmonella LT, and UPEC CFT073) within THP-1 macrophages over 24 hours. D) Imaging of intracellular bacterial communities (IBCs) within Lamp1-positive vacuoles in THP-1 macrophages by AIEC, commensal, and pathogenic strains at 24 hours post-infection (P.I.). The scale bars are 5 µm.

### In depth analysis of the genome content of five different AIECs

To refine our understanding of macrophage colonization, we curated a subset of four new AIECs spanning the B1, B2, and A phylogroups. Employing a combination of Illumina and Nanopore sequencing techniques, we assembled the complete genomes of each strain (Fig. 2A and Supplementary Table S1 and S2, see Methods). Notably, we achieved a single contig for the chromosome of each strain and identified several plasmids ranging from 1.5 kb to 133 kb (as illustrated in Supplementary Fig. S2A and S2B). Large plasmids found in strains CEA601S, CEA212U, CEA224S, and CEA106S exhibit significant homologies with the NRG857c and UM146 plasmids (Supplementary Fig. S2C), yet show no similarity with the Cyrano phage-plasmid from LF82 (Miquel S et al., 2010; Misson P et al., 2023; Nash JH et al., 2010). Even within this limited bacterial cohort, genomic diversity remains striking, with nearly one-third of the genome comprising shell or singleton genes (Prophages, phage defense systems, toxin-antitoxin systems, and metabolic islands exhibit notable disparities (Supplementary Fig. S3). None of the prophages hosted by LF82 were identified in the four other strains. Instead, they possess different complete or incomplete prophages, possibly inserted at analogous genomic loci. While some of these prophages may carry virulence factors, such as the Shiga toxin in PhiP27-like prophage or GtrII-GtrA proteins in phage SfII, meticulous examination revealed that none of these virulence factors were actually encoded by the prophages within our study group (see Supplementary Table S2).

**Figure 2:**
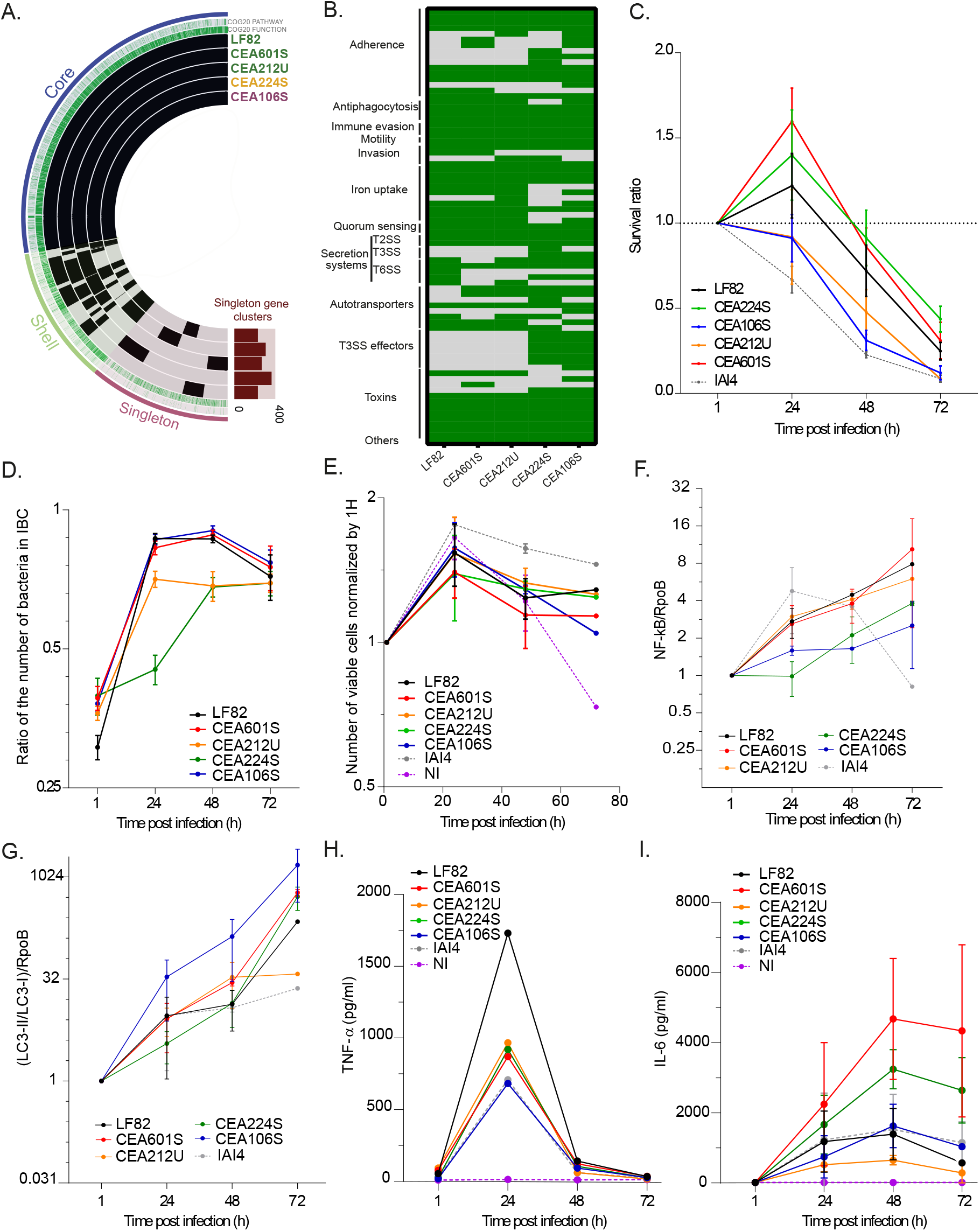
Analysis of virulence arsenal and macrophage response to five AIEC. A) Presence/absence chart of putative virulence factors in AIEC genomes. The layers represent individual genomes organized by their phylogenomic relationships. In the layers, black indicates the presence of a gene cluster, while light colors indicate its absence. Detailed graphs are presented in Supplementary Figure 3. B) Correlation matrix of the virulence arsenal across five AIEC strains. C) Survival of AIEC strains within macrophages during a 72-hour infection kinetics study. Error bars represent the mean and standard error of the mean (SEM); n ≥ 3. Individual data points are shown in Supplementary Figure 7B for each strain and Supplementary Figure 7C for each time point. Data were analyzed using the Mann-Whitney test: *P < 0.05, **P < 0.01, ***P < 0.001, ****P < 0.0001. D) Quantification of bacteria that form intracellular bacterial communities (IBCs) relative to the total bacterial population. Error bars represent the mean and SEM. E) MTS assay of THP-1 macrophages during the infection kinetics with AIEC strains. Error bars represent the mean and SEM. F) Inflammatory response of infected macrophages, measured by NF-κB levels normalized to RpoB and 1-hour post- infection (P.I.). Error bars represent the mean and SEM. G) Autophagy response of infected macrophages, indicated by the LC3II/LC3I ratio normalized to RpoB and 1-hour P.I. Error bars represent the mean and SEM. H) TNF-α quantification in the culture media of macrophages infected by AIEC or the commensal strain IAI4. I) IL-6 quantification in the culture media of macrophages infected by AIEC or the commensal strain IAI4. Error bars represent the mean and standard deviation (SD).

The four newly identified AIECs manifest highly diverse pathogenicity gene profiles (Fig. 2B and Supplementary Fig. S3), potentially contributing to distinct strategies for macrophage colonization. Even strains from the B2 group (CEA601S and CEA212U) exhibit divergent sets of fimbriae, invasion factors, agglutinins, colicins, iron capture systems, and secretion systems compared to LF82. This suggests that LF82’s colonization strategy may not be universally adopted among other members of our study group.

#### Secretion

The secretion of virulence factors has been extensively shown to influence the colonization of host cells by intracellular pathogens (Poirier V & Av-Gay Y, 2015). Research on pathogen models such as *Salmonella*, *Shigella*, and *Yersinia* has elucidated the essential roles played by Type 3 secretion systems (T3SS). In addition to the general Type 2 secretion system, the LF82 genome exhibits two complete and one partial Type 6 secretion systems (T6SS), a T4SS, and numerous autotransporters (T5SS), but lacks a T3SS (Fig. 2B and Supplementary Fig. S3B). However, the involvement of these systems in the colonization of epithelial cells or macrophages by LF82 has not yet been observed.

Regarding secretion, strain CEA224S (B1 group) stands out as the most distinctive, as it harbours a complete type III secretion system on its large plasmid, suggesting its capability to secrete virulence factors into its host (Supplementary Fig. S3B). Furthermore, it also carries a partial type III secretion system similar to the PrgK, PrgI, and OrgB proteins located between the *yqeG* and *ygeR* genes on the chromosome (Supplementary Table S2). This cluster is also present in strain CEA106S (A group) at the same position. LF82 exhibits two complete type VI secretion systems, defined as pathogenicity islands PAI-I and PAI-III (Miquel S et al., 2010). Strains CEA212U, CEA224S, and CEA106S also contain a T6SS at the PAI-I location, while strain CEA601S harbours a large gene cluster encoding sugar metabolism enzymes, transporters, and transcription factors in this position. Strains CEA212U and CEA224S also display putative T6SS at the PAI-III locus, whereas this locus is empty in strains CEA601S and CEA106S. All strains exhibit a wide array of putative autotransporters (T5SS) and one or two T4SS frequently found on their respective plasmids (Supplementary Fig. S3B).

#### Appendages

AIECs exhibit various appendages that may facilitate motility, adherence, invasion, and protection from phagocytosis (Fig. 2B). Motility is mediated by the flagellum, which is present in every strain. Adherence is facilitated by fimbriae, which can vary in type. All strains possess the operon necessary for the synthesis of type 1 fimbriae (*fimA-H*). Additionally, LF82 harbors a gene cluster for long polar fimbriae, while CEA601S and CEA106S feature afimbrial adhesins (*afaABCD*), CEA212U contains a P fimbriae gene cluster (*papA-G*), and CEA224S possesses both adhesive fimbriae (*faeC-J*) and long polar fimbriae (Supplementary Fig. S3A). Each AIEC strain encodes capsule synthesis gene clusters. In LF82, capsule synthesis is enabled by the PAI-IV. This locus exhibits high polymorphism; it consistently contains genes involved in capsule synthesis and a putative secretion system as in LF82, but may also contain genes for the P fimbriae gene cluster (*papA-G* operon), phosphoglycerate transport (*pgtP-pgtCBA*), or virulence factors (*sat* protease, colicin, *espK*) in other strains. The *E. coli* common pilus (ECP) gene cluster is present in all strains, promoting binding to the extracellular matrix (Mondal R et al., 2022); CEA212U and CEA106S additionally display FdeC, a homolog of Invasin and Intimin, which facilitates kidney and bladder colonization by UPEC (Nesta B et al., 2012). Other invasion factors such as Tia/Hek, IbeABC (Cieza RJ et al., 2015), and Hra were detected in these strains.

#### Iron capture

Iron acquisition is essential for bacteria as a nutritional requirement. As part of the nutritional immunity response host secrete sequestration proteins to deplete free iron and limit pathogen growth (Ullah I & Lang M, 2023). AIEC genomes contain more iron capture systems compared to other *E. coli* (Dogan B et al., 2014). We demonstrated that activation of the Yersiniabactin production island HPI (PAI-II) in LF82 corresponds to the beginning of bacterial multiplication within macrophages. This prompt us to investigate the distribution of iron capture systems in our bacterial cohort (Supplementary Fig. S3D). The PAI-II locus is occupied by the HPI in LF82, CEA601S, CEA212U and CEA106S but empty in CEA224S. Strains from the B2 group also display the Chu haeme capture system.

#### Virulence effectors

The genomes of AIEC encode several potential offensive virulence factors (Supplementary Fig. S3A). These include Vat and Hbp proteases in LF82, CEA601S, and CEA212U; Colicins in CEA212U; and Sat and SenB toxins in CEA106S. Strains CEA224S and CEA106S, which possess putative T3SS, also carry the same group of secreted factors: EspX5, EspX1, EspR1, EspX4, EspL4, and EspY1. Among these, only EspY1 has known host cell targets related to apoptosis or cell cycle regulation.

This diverse arsenal suggests that these bacteria may react differently in various host environments to colonize tissues or cells. This highlights the importance of conducting thorough phenotypic analyses before designing any potential therapeutic strategies. Furthermore, this necessity is underscored by the significant antibiotic resistance potential observed in our AIEC cohort (Supplementary Fig. S3C).

### Virulence of AIECs within macrophages

We assessed bacterial viability over a 72-hour period post-infection. An equal amount of exponentially growing AIEC strains and IAI4 commensal strain were exposed to THP1 macrophages differentiated from monocytes for 20 minutes. This resulted in an infection rate of 1 to 10 bacteria per macrophage for LF82, CEA601S, CEA212U, and CEA106S, slightly higher rate observed for CEA224S 1 hour after infection (with 88% of macrophages infected, including 12% with IBCs) and LF82 (with 85% of macrophages infected but only 8% with IBCs) compared with the other strains (between 46% and 62% of macrophages infected and between 5 and 8% with IBCs, Supplementary Fig. S1B). For LF82, CEA224S, and CEA601S, the proportion of viable bacteria remained relatively stable for 48 hours before decreasing to 0.3 - 0.5 at 72 hours (Fig. 2C). In contrast, the numbers of CEA106S and CEA212U declined more rapidly, suggesting less efficient survival, evasion from macrophages, or more frequent macrophage killing.

Throughout this infection kinetics, the majority of AIEC bacteria resided within Intracellular Bacterial Communities (IBC), ranging from 60 to 94% between 24 and 72 hour post-infection depending on the strain (Fig. 2D and Supplementary Fig. S1B). Even strains with low colony-forming units (CFU) at 72 hours, such as CEA106S and CEA212U, exhibited IBC frequencies exceeding 70% (Fig. 2D). These IBCs started as small communities (< 10 bacteria) at 1 hour and increased in size, reaching a maximum of 392 bacteria per one IBC for CEA106S at 48h post-infection (median = 22.6; IQR = 38.5; Range = 389.2) (Supplementary Fig. S1C). Thus, IBC appears to be the preferred lifestyle of AIECs within macrophages despite variability between genomes.

To investigate the impact of IBC on macrophages, we assessed metabolic activity, cell viability, inflammation, autophagy, and cytokine production. We utilized formazan to monitor ATP synthesis in infected macrophages (Fig. 2E), which remained constant throughout the infection and comparable to non-infected controls. This suggests that macrophage viability was not significantly affected by the infections and therefore not explaining the decline in bacterial population at the 72 hour mark. We quantified NFκB, a marker of inflammation, and LC3-II, a marker of autophagy. Consistent with existing literature, strains LF82, CEA601S, CEA212U, and our commensal control IAI4 induced inflammation (Fig. 2F and 2G), with approximately 4 times more NFκB in macrophages at 48 hour post-infection. Surprisingly, strains CEA106S and CEA224S did not induce inflammation, respectively we observe 1.6 and 2 times more NFκB at 48 hour post infection. Additionally, CEA601S, CEA212U, CEA224S, and CEA106S promptly induced autophagy (respectively 28, 34, 14, 135 times more LC3-II conversion at 48 hour than at 1hour post-infection), whereas LF82 did so latter during infection (13 times more at 48 hour post infection and increased considerably to 223 times at 72 hour post infection like CEA224S, CEA601S and CEA106S) (Fig. 2F and 2G). We measured cytokine production by the infected macrophages and the presence of bacteria in macrophages promotes the secretion of TNF-α compared to uninfected macrophages at 24 hours post-infection. TNF-α levels being twice as high in macrophages infected by LF82 (1700 pg/mL of TNF-α). However, TNF-α levels significantly decrease at 24 hours for all strains (Fig. 2H). Measurement of IL-6 production by macrophages shows a high amount of IL-6 for the CEA601S and CEA224S strains, with an increase over time (Fig. 2I). Other AIEC strains promote less IL-6 secretion, but it remains sustained over time.

### Adaptation of AIECs within macrophages

We hypothesized that variation of macrophage responses may reflect different AIEC adaptation strategies. We recently described that within IBC, LF82 expressed genes from the biofilm pathway were labelled with the SBA and WGA lectins and anti-curli antibody. Moreover, FRAP experiments showed that bacteria within IBC are completely static. These observations suggested that IBC were structured by a matrix and could be considered as micro-biofilm structures (Prudent V et al., 2021). Surprisingly, only LF82 and CEA106S exhibit strong WGA labelling. In contrast, the strain CEA224S exhibits a weak WGA labelling visible in some of the IBC and strains CEA601S and CEA212U did not exhibit any WGA labelling (Fig. 3A). This is in good agreement with different movements of the bacteria within IBC (Supplementary Fig. S4). IBC are therefore of different nature, AIECs either multiply within biofilm-like structures or less rigid environments where only adherence between bacteria and between bacteria and the phagolysosomal membrane may limits their movements. Another important factor contributing to macrophage colonization by LF82 is the ability to switch from a replicative to a non-replicative state and therefore to stimulate the appearance of antibiotic tolerant persisters (Demarre G et al., 2019). As proxy of replicative status, we measured persisters frequency at 1, 24, 48 and 72 hours P.I. for each strain. CEA601S and CEA224S followed the same trend as the one observed for LF82, persisters frequency increased during infection, reaching ⁓ 20% of the population. By contrast, CEA212U and CEA106S followed IAI4 trend, exhibiting very low and nearly constant level of persisters (Fig. 3B). The acidic environment induces *Salmonella*’s persisters; we tested the acid response of the five strains with a biosensor based on the *asr* promoter fused with an unstable GFP. Flow cytometry analysis of lysed macrophages revealed two types of behaviours: strains LF82, CEA601S, CEA224S and IAI4 exhibit a frequent and strong (Fig.3C and Supplementary Fig. S5B) expression of the *asr* promoter; by contrast strains CEA212U and CEA106S rarely exhibit expression of *asr* promoter (Fig. 3C). The findings corroborate with the *asr* expression results. At 24 hours post-infection, CEA106S and CEA212U have lower *asr* gene transcription compared to other AIEC strains (Fig. 3D). This suggests these strains are in a less acidic environment. Macrophage acidification measurements from 1.5 to 24 hours indicate that CEA106S resides in macrophages in a less acidic environment than LF82, similar to non-infected ones (Fig. 3E). Microscopy confirmed these observations, IBC formed by LF82, CEA601S and CEA224S frequently exhibit *asr*+ bacteria while those formed by CEA212U and CEA106S did not contain *asr*+ bacteria (Supplementary Fig. S5A). These images confirmed our previous LF82 observations, showing that within IBC, bacteria respond heterogeneously to the acid pH stress. The good correlation between persisters frequency and *asr* induction prompted us to check if multiplication rate differed in these two groups. We used a biosensor constructed with the rrnBP1 promoter to monitor stringent response and a FtsZ-GFP ring reporter for cell division. Concerning stringent response, the five AIECs and IAI4 behave similarly, only a small proportion of the bacteria (5 – 15%) exhibit high *rrnBP1* expression and therefore high translation capacity (Supplementary Fig. S6A). We conclude that most of them encounter nutrient depletion and engage stringent response. Since the IAI4 bacteria that do not survive within IBC show comparable level, we conclude reduction of *rrnBP1* expression rather reveal *E. coli* adaptation to the poor nutritive environment than AIEC adaptation. LF82 and CEA106S presented twice as much bacteria with FtsZ ring at 48h (respectively 29% and 27%) compared to 24h (respectively 17% and 24%) while similar number of dividing cells (30%) where observed for CEA212U and CEA224S at both timing (Supplementary Fig. S6B and S6C). Unfortunately, we were not able to collect FtsZ data for CEA601S. In controlled conditions, the frequency of FtsZ ring formation varies with growth rate and the frequencies observed within IBC would corresponds to a two hours generation time (Den Blaauwen T et al., 1999). However, considering the significant heterogeneity of the bacterial population inside macrophages and the influence of stresses such as acid (Peters K et al., 2016) or micro-aerobiosis (Murashko ON & Lin-Chao S, 2017) the frequency of FtsZ may not directly correlate with growth rate as it does in test tube conditions. This relationship between FtsZ ring assembly and multiplication is particularly altered for the strain CEA212U that form giant filaments at late time points post infection (Supplementary Fig. S6D). This phenotype, comparable to UPEC filamentation in bladder cells, suggests either that CEA212U encounters a different environment compared to other AIEC or express a particular virulence factor.

**Figure 3:**
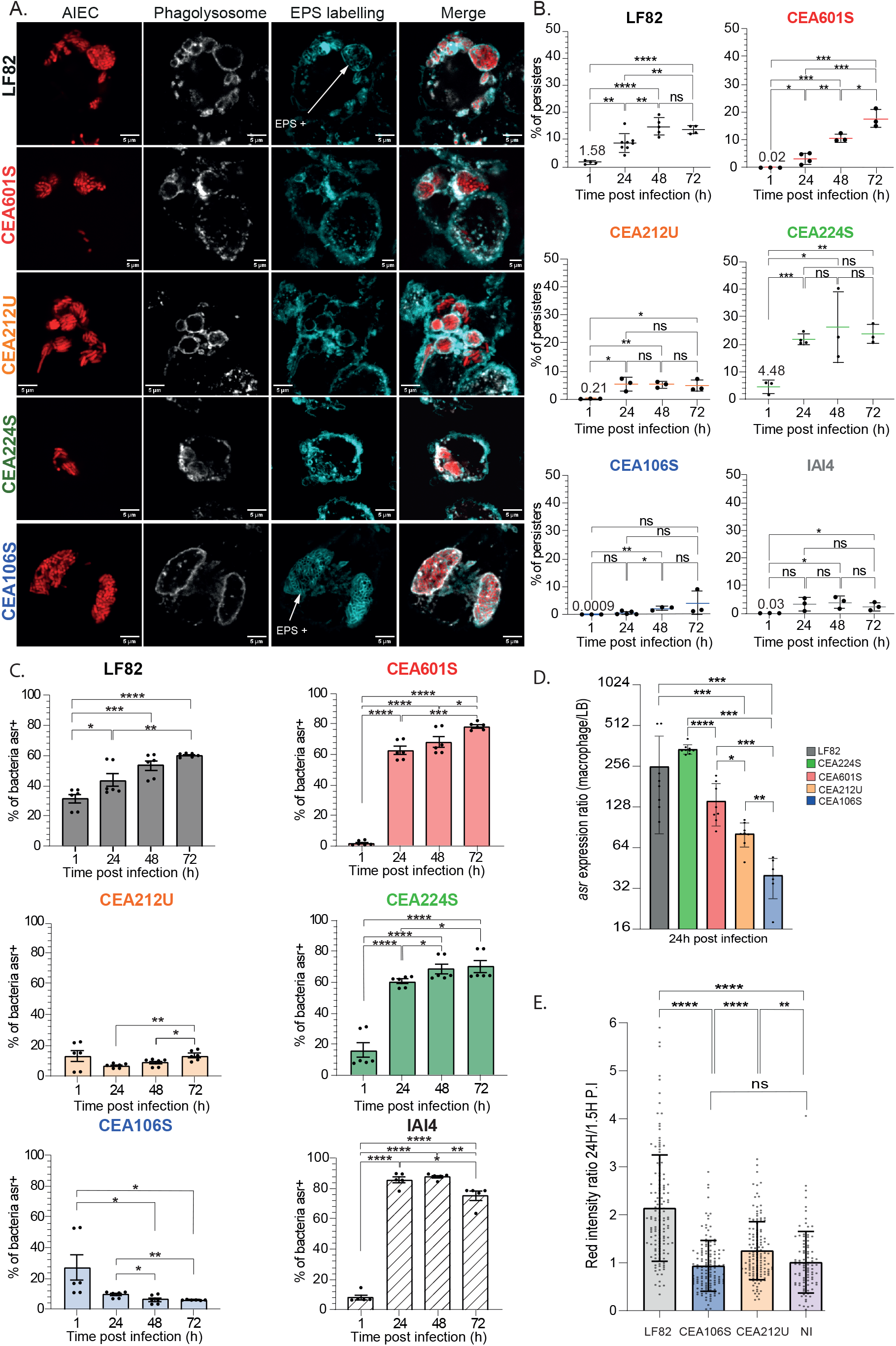
Characteristics of IBC forms by different AIECs. A) Images of IBCs formed by AIEC, showing WGA lectin-labeled polysaccharides and phagolysosomes marked by Lamp-I. Arrows indicate WGA labeling, suggesting the presence of an intracellular polysaccharide matrix. Scale bars represent 5 µm. B) Measurement of persister frequency during AIEC growth within macrophages. P-values are indicated (Student’s t-test: *P < 0.05, **P < 0.01, ***P < 0.001, ****P < 0.0001); error bars represent the mean and standard deviation (SD); n ≥ 3. C) Flow cytometry quantification of AIEC response to intracellular acid stress (asr promotor). P-values are indicated (Student’s t-test: *P < 0.05, **P < 0.01, ***P < 0.001, ****P < 0.0001); error bars represent the mean and standard error of the mean (SEM); n ≥ 3. D) qRT-PCR analysis of AIEC acid stress response induction at 24 hours post-infection (P.I.) in THP-1 macrophages compared to the stationary phase in rich liquid medium. Data represent relative quantification of asr, normalized by the *rpsM* reference gene. P-values are indicated (Mann-Whitney test: *P < 0.05, **P < 0.01, ***P < 0.001, ****P < 0.0001); error bars represent the mean and standard deviation (SD); n ≥ 3. E) Quantification of pHi probe intensity in macrophages infected by AIEC strains LF82, CEA106S, CEA212U, or uninfected. Each point represents the intensity ratio at 24 hours post-infection compared to the mean intensity at 1.5 hours. P-values are indicated (Mann-Whitney test: *P < 0.05, **P < 0.01, ***P < 0.001, ****P < 0.0001); error bars represent the mean and standard deviation (SD); n ≥ 3.

Altogether, these results suggest that within IBC the average translation capacities of each strain are low, nevertheless, an important number of bacteria manage to multiply or elongate (CEA212U) during the whole infection. AIEC strains, however, differ significantly in the frequency of persisters generated over time (Supplementary Fig. S7A) and these differences correlate well with their induction of the acidic response. Finally, we observed the formation of a strong exopolysaccharide labelling within LF82 and CEA106S IBCs, these two strains also exhibit an increase in the frequency of FtsZ ring over time, suggesting that the EPS matrix may contribute to stimulate proliferation within IBC for these strains.

### Fates of AIECs and infected macrophages

The dissemination of AIEC outside macrophages has not yet been documented. To assess whether colonization of macrophages by AIEC represents a dead end that may not benefit the bacteria or a springboard for infection propagation, we monitored infected macrophages for 12 to 16 hours at the 48-hour time point. Frequent imaging at low laser intensity enabled us to track moving macrophages accurately, detect rapid events while preserving macrophage and bacteria viability. We observed hundreds of infected macrophages for each strain (Fig. 4A-F). Outside dissemination was observed for every strain; in most cases, it was concomitant with macrophage death. However, in the case of LF82, CEA212U, and CEA106S, we also observed vacuolar rupture and dissemination within the cytosol of the macrophages. Additionally, every strain exhibited vacuole fusions, with these fusions being particularly frequent with CEA224S. More surprisingly, we observed the transfer of intact IBC from one macrophage to another for LF82, CEA601S, and CEA224S; most of the time this transfer occurred concomitantly with the death of the donor macrophage but it was also observed between live macrophages, resembling a process similar to vomocytosis. For CEA212U, at 48 hours P.I. many filaments were observed, in the following hours they either continue to grow or resolve forming clonal IBC. Therefore, CEA212U filamentation did not directly facilitate dissemination as it does for UPEC in bladder cells. We suspect, therefore, that filamentation is part of a response to a transient stress appearing in the vacuole. It is noteworthy that inter-macrophage transfers occurred in the presence of gentamicin in the medium, which did not seem to affect IBC integrity. We measured the viability of released bacteria in a two-hour window without gentamicin. LF82 and CEA601S exhibited a release rate of 1% of the total population in 2 hours, while CEA106S exhibited a much higher release rate of 10% in two hours (Fig. 4H). Optical density measurements confirmed that this difference could not be explained by bacterial multiplication, which only started 4 to 8 hours after release (Fig. 4I). These observations unambiguously demonstrate that dissemination is possible for all strains and therefore that IBC is not a dead end for AIECs.

**Figure 4:**
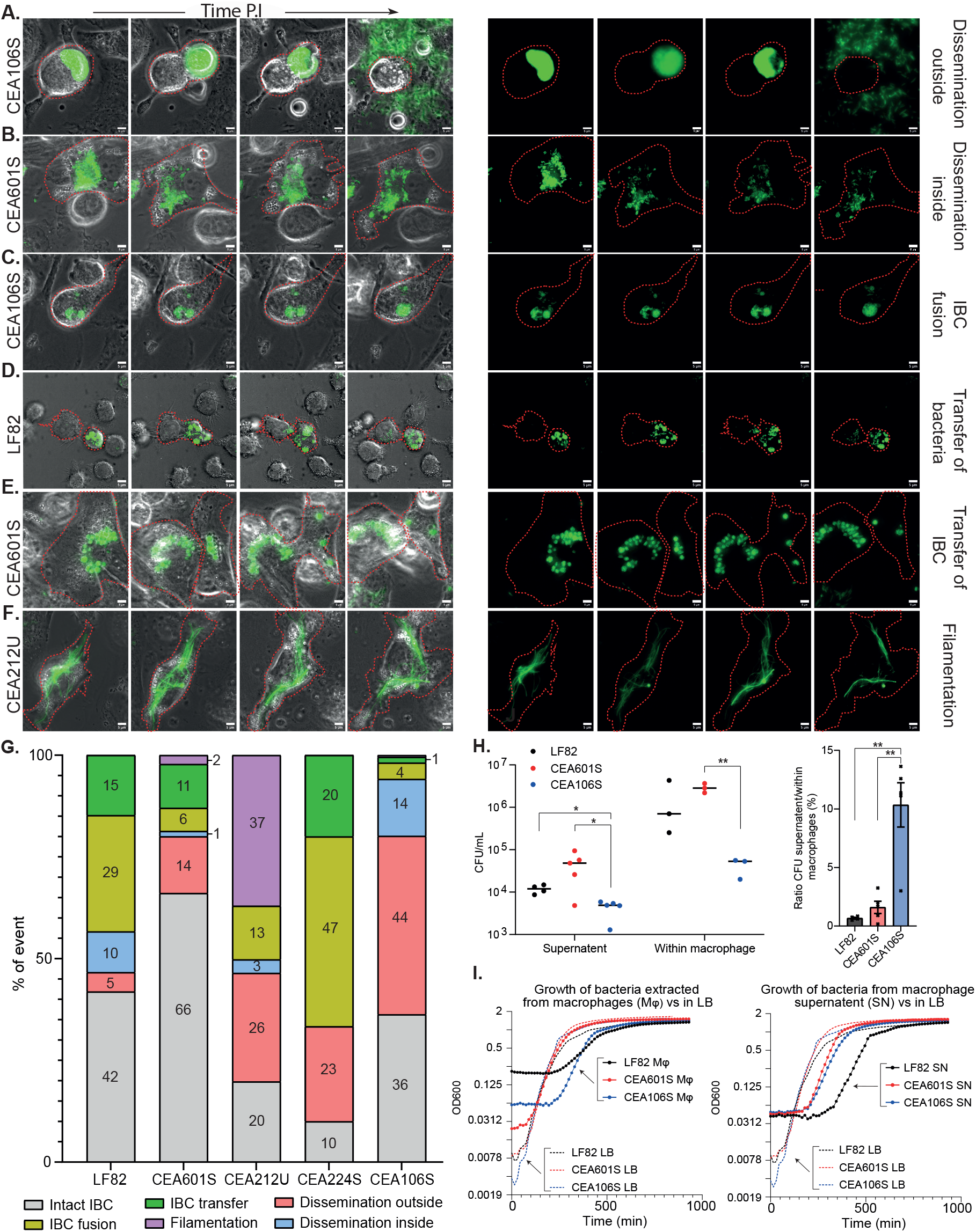
Fates of AIECs and infected macrophages. Live imaging assembly of THP-1 macrophages infected by AIEC, with recording starting at 48 hours post-infection (P.I.). Bacteria are shown in green, and macrophages are outlined in red. Scale bars represent 5 µm. A) Dissemination of IBC outside macrophages. B) Dissemination of IBC within a macrophage. C) IBC fusion. D) Transfer of an IBC to another macrophage, resulting in rupture of the recipient cell. E) Transfer of an IBC from a donor to a recipient macrophage. F) AIEC filamentation. G) Quantification of different IBC fates within THP-1 macrophages. H) Proportion of viable bacteria at 24 hours P.I. in the supernatant of macrophages or within macrophages. The graph on the right represents the ratio of bacteria in the supernatant to bacteria within the macrophages. P-values are indicated (Student’s t-test: *P < 0.05, **P < 0.01, ***P < 0.001, ****P < 0.0001); n ≥ 3. I) Recovery of bacterial growth from within macrophages (left) or from macrophage supernatant (right), compared to growth in LB medium (dashed line).

### Correlations between AIEC phenotypes, genomic contents and pathogenicity

In the course of this study, we collected about 60 phenotypic characteristics for the five AIEC strains that helped us to decipher macrophage colonization and dissemination properties. To identify key phenotypes that would allow screening other AIECs with less effort, we quantitatively analyzed the correlation of these phenotypes with the AIEC survival (CFU). Three particular phenotypes strikingly correlate with survival: the production of IL-6 at 24h (R^2^ > 0.85), the frequency of persisters at 48h P.I. (R^2^ > 0.8) and the acid response of the bacteria at 24h P.I. (R^2^ > 0.9) (Figure 5A-C). Suggesting that these three characteristics are connected. The link between acidity of the vacuole and persisters was documented for *Salmonella* infection (Helaine S et al., 2014). By contrast, the connection between IL-6 production and phagosome acidification or persister formation is not yet documented. These observations suggest that AIEC that deal positively with macrophage pro-inflammatory response (M1 state) colonize them more efficiently. Surprisingly we did not observe similar correlation with NF- and TNF-α production. Although more strains will be required to validate these observations, they open the way to simpler screening of AIEC collections.

**Figure 5:**
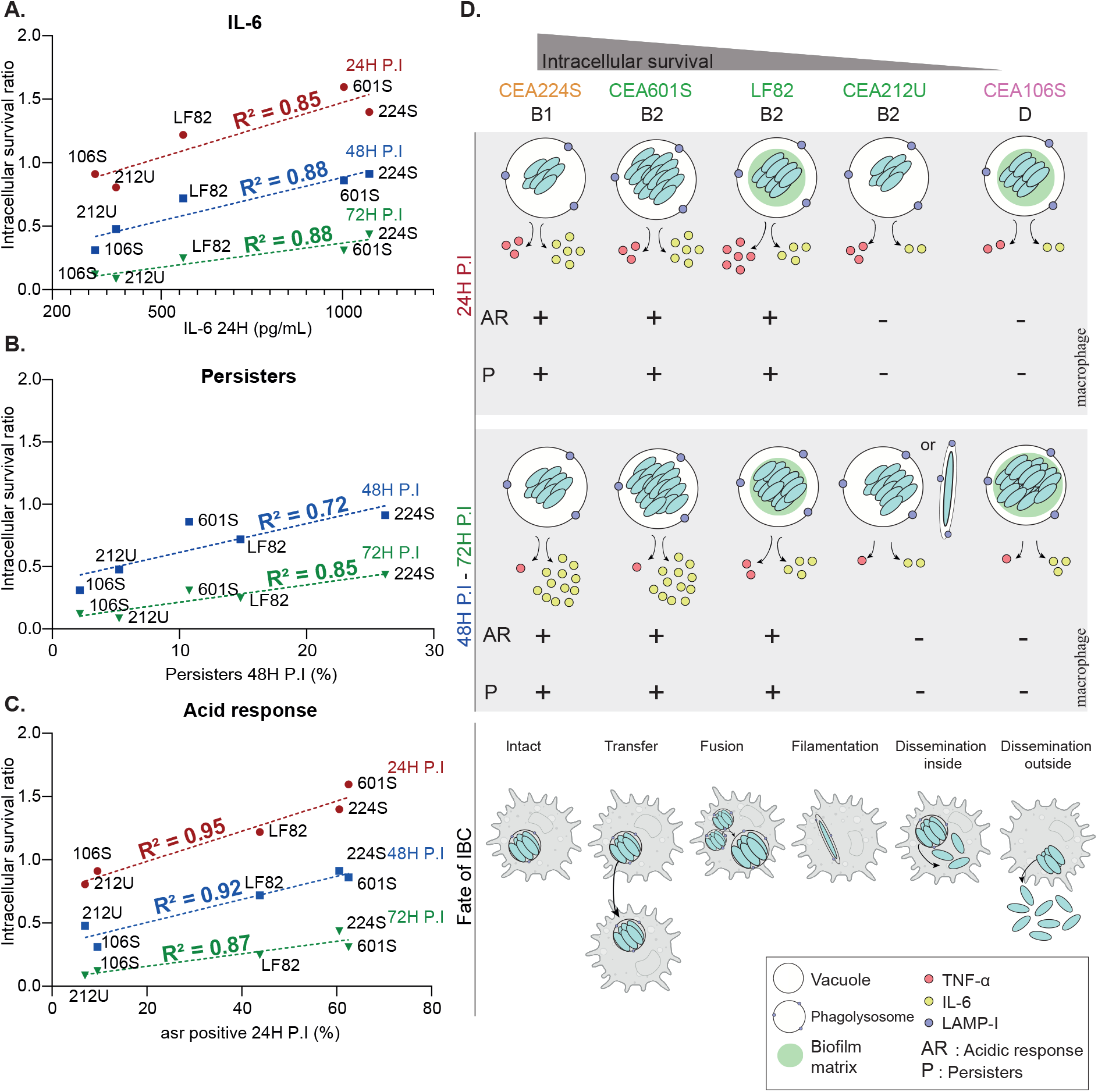
IBC formation relies on different strategies. Pearson correlation between intracellular survival and: A) IL-6 production secreted by infected macrophages; B) formation of persisters; C) acidic response of AIEC. The red dashed line represents the correlation at 24 hours post-infection, the blue dashed line at 48 hours, and the green dashed line at 72 hours. The correlation coefficient is indicated by R². D) Overview diagram of the various survival strategies employed by the five AIEC strains studied, along with their outcomes.

## Discussion

### Convergent evolution of AIECs leads to IBC formation

Alterations in the composition and function of gut microbial communities are implicated in inflammatory bowel disease (IBD), with Crohn’s disease (CD) patient microbiomes showing an increase in mucosa-associated Enterobacteriaceae, including AIECs (Baumgart M et al., 2007; Glasser AL et al., 2001). AIEC are defined by their *in vitro* adherent-invasive behavior. There is a clear need to extend our knowledge of AIEC to pathoadaptive determinants of colonization and virulence. Our study of 10 new AIEC strains confirmed previous results indicating that the AIEC phenotype encompasses *E. coli* strains with very different genome contents (Camprubí-Font C et al., 2018; Dogan B et al., 2014; Wang J et al., 2021). This genomic diversity is attributed to both SNP polymorphisms of core genes (Fig. 1C) and horizontal gene transfer (Fig. 1B). AIECs in our collection belong to the A, B1, B2, and D phylogroups. Despite this diversity, all tested AIECs exhibited a common phenotype: the formation of intracellular bacterial communities (IBC) within Lamp-1 positive compartments inside macrophages. These IBCs differ significantly from those of *Salmonella*, *Shigella*, or uropathogenic *E. coli* (UPEC) observed under the same conditions (Fig. 1). To refine this genomic-phenotypic analysis, we assembled and annotated the genomes of four new AIEC strains (CEA106S, CEA212U, CEA224S, and CEA601S). Compared to LF82, each of these strains carries over 350 specific genes, with only 31 genes shared among the four and LF82 but absent from a commensal control. The genomes of AIECs are particularly rich in prophages, which may carry morons (Misson P et al., 2023; Supplementary Table S2), and they also contain conjugative plasmids that carry virulence factors such as T3SS in CEA224S. Each studied AIEC harbor a distinct set of putative virulence factors, including siderophores, adhesins, effectors, toxins, and secreted proteases (Supplementary Fig. S3). This rich genomic diversity may result from fortuitous evolution (Diard M & Hardt W-D, 2017) and various selective pressures that AIECs encounter in their diverse niches (Touchon M et al., 2020; Wylie KM et al., 2019). Therefore, we propose that convergent evolution leads to the IBC phenotype. This suggests that in CD patients, macrophages or other immune cells impose selective pressure on mucosa-associated Enterobacteriaceae. Supporting this hypothesis, the colonization of *lamina propria*’s immune cells by proteobacteria during CD (Dheer R et al., 2020) and the strong positive correlation between AIEC replication in macrophages and the severity of both macroscopic and microscopic lesions in mice (Kittana H et al., 2023) have been observed.

### IBC formation relies on different strategies

We observed that AIECs relies on different strategies to build IBC (Fig. 5).

**LF82, CEA601S and CEA224S**: These three strains display the best survival rate at early and late time point of the infection (Fig. 2C). Their IBC grow during the first 48h in an acidic phagolysosome and they continuously produce more persisters. This pattern follows the earlier characterization of LF82 colonization (Demarre G et al., 2019; Prudent V et al., 2021). CEA224S, from the B1 phylogroup, presents a less intense WGA labelling as observed for LF82 and CEA106S (Fig. 3A). By contrast, CEA601S, from B2 phylogroup, did not exhibit WGA labelling and individual bacteria are more mobile with IBC (Supplementary Fig. S4). These observations suggest that a robust adhesive matrix is not required for IBC formation and bacterial survival. This might also be the case for IBC formed by UPEC in bladder cells (Zhou Y et al., 2023).

**CEA106S:** At first glance, our CFUs and IBC counting data suggested that CEA106S is the less efficient AIEC of our group for macrophage colonization (Fig. 2C). These observations might actually be misleading because dissemination is very frequent with this strain. Bacteria grow efficiently (high frequency of bacteria with FtsZ ring, important EPS staining and low amount of persisters), they form large IBC, that reach carrying capacity and explode liberating live bacteria in the medium that are not counted by a classical gentamycin protection assay. In good agreement with the hypothesis of an efficient multiplication, we observed that although inflammation and autophagy were activated during CEA106S infection, the bacteria stopped rapidly the induction of its acid stress response (Fig. 3C and Supplementary Fig. S5A and S5B). Altogether, these observations suggested that CEA106S is able to detoxify its vacuole (Fig. 3E). Considering its filiation with the A phylogroup where a majority of commensal *E. coli* were identified this is particularly surprising.

**CEA212U:** CEA212U belongs to the canonical B2 phylogroup, however it behaves closely to CEA106S from the A phylogroup. It did not produce high level CFU at 48 and 72 hours (Fig. 2C), did not induce acid stress and did not accumulate persisters (Fig. 3B and 3C). Strikingly, it elongates forming large filaments that may utterly resume cytokinesis. This suggest that they are metabolically active. Although the filamentation phenotype recall from UPEC dissemination strategy, we do not think CEA212U uses it as a dissemination strategy. Dissemination is only observed for IBC containing small cells (Fig. 4). Moreover, genomic comparison did not reveal common gene sets shared with UPEC and missing from other AIECs that could explain this phenotype.

### Dissemination of AIECs

Earlier studies on macrophage colonization by LF82 neither escapes from the phagolysosome nor induces macrophage death (Glasser AL et al., 2001). Moreover, RNA-seq analysis indicated that at early time points, LF82 induces anti-apoptotic functions in macrophages (Prudent V et al., 2021). This led to the general consensus that AIEC replicate within macrophages without causing host cell death (Yao T et al., 2023). However, the results presented sharply contradict this view (Fig. 4). Our findings indicate that while AIECs do disseminate to the external medium and synchronously kill the macrophage, they also spread by other means. Some strains break the vacuolar membrane and disseminate into the cytosol, while others disseminate as an intracellular bacterial community (IBC) that can be recaptured by other macrophages, subsequently fusing with their endogenous IBC. Additionally, we observed events similar to the vomocytosis process, a transfer of loaded phagolysosomes from a live macrophage to a new macrophage, a phenomenon described for fungal pathogens (Johnston SA & May RC, 2013). Interestingly, the frequency of these dissemination processes in the population of infected macrophages varies from one strain to another. We postulate that these differences may reflect specific macrophage-pathogen interactions.

### Could IBC characteristics be used as a marker of AIEC pathogenicity?

The search for genomic markers of the AIEC phenotype was unsuccessful. Given that bacterial survival within immune cells has been identified as a marker of Crohn’s Disease (CD) (Dheer R et al., 2020) and a marker of macroscopic and microscopic intestinal lesions in mice (Kittana H et al., 2023), we hypothesized that an in-depth analysis of the phenotypes associated with AIEC survival within macrophages would help classify them. First, we demonstrated that all AIECs, unlike *Salmonella*, *Shigella*, UPEC, or commensals, form IBC. Therefore, the presence of IBCs inside macrophages differentiates AIECs from other enterobacteria but does not distinguish between different AIEC strains. The analysis of more than 60 phenotypic characteristics of AIEC infection allowed us to identify some that are interconnected. For instance, intracellular survival, persister frequency, vacuole acidification, and IL-6 production were highly correlated. In contrast, the WGA labelling of a putative extracellular matrix, which appeared to reinforce LF82 survival (Prudent V et al., 2021), did not correlate with the intracellular survival of other strains. This suggests that successful AIECs, those with higher survival rates, rely more on their ability to cope with macrophage responses rather than on their ability to build a protective matrix. This observation generalizes to the AIEC group previous findings showing that acidification and inflammation play key roles in LF82 multiplication (Bringer M-A et al., 2006). We add to this scenario the significant contribution of continuous persister production during growth under stress. Although additional validations with more strains and *in vivo* assays are required, we propose a minimal model for optimal AIEC multiplication. It necessitates phagolysosomal activation for multiplication and simultaneous persister formation. IBCs formed in this toxic environment will accumulate induced persisters that tolerate stress, while replicating bacteria should be at risk, and their debris will stimulate macrophage signalling, including IL-6 production. IL-6 would favour the recruitment of other immune cells to the infection site, potentially aiding the dissemination of AIECs to other immune cells (Fig. 5D). Additional options such as extracellular matrix production might give an extra advantage to some strains. Alternatively, some strains may select a different path involving phagolysosome detoxification that appeared less efficient, possibly because it does not involve IL-6 production. IL-6 imbalance has been documented in CD, and it is proposed that it favours an intestinal microenvironment prone to chronic inflammation. Thus, targeting IL-6 could be part of therapeutic strategies (Alhendi A & Naser SA, 2023).

## Methods

### Strains and plasmids

Strains and plasmids are described in Table S3. The collection of new *E. coli* AIEC strains (Buisson A et al., 2021) was sequenced by Illumina technology to generate draft genome contigs. For the strains CEA212U, CEA106S, CEA224S and CEA601S genomic DNA was extracted with Invitrogen PureLink. Total DNA was sequenced by Nanopore technology (Novogen platform). To achieve *de novo* hybrid assembly of short and long reads passing quality control and filtering, we used Unicycler v0.5.0 tool, with default parameters for chromosomes and plasmids (Wick RR et al., 2017).

### Genome analysis and plasmid content

The genome annotation was performed with PROKKA v1.14.6 and based on LF82 annotation (Seemann, 2014). *E. coli* phylogroup was determined with ClermonTyping (Beghain J et al., 2018; Clermont O et al., 2019) and serotype with ECTyper (Bessonov K et al., 2021).Circos on plasmids sequence was performed using Galaxy (Krzywinski M et al., 2009; Rasche H & Hiltemann S, 2020). To identify antimicrobial resistance genes in chromosomes and plasmids we used AMRFinderPlus version 3.11.26 with database version 2024-01-31.1. To search virulence factors, we used VFDB on fasta assembly files, for TSS and defense systems we used MacSyFinder and DefenseFinder/CRISPRCasFinder in MaGe (Vallenet D, 2006; Vallenet D et al., 2019).For prophage we used Phaster and then Phigaro in MaGe. Finally, identification of T/AT systems was carried out with TAfinder2.0 (Xie Y et al., 2018).

### Pangenome analysis

To characterise the distribution of the gene content, comparative analysis of the genomes of fifteen *E .coli* was performed through the pangenomics pipeline Anvi’o v7.1 (Eren AM et al., 2015). The resulting pangenome was visualised in an interactive interface by running the script anvi-display-pan, where strains are clustered based on gene frequencies and the phylogram is clustered based on presence/absence of pattern of gene clusters. The Pan-genome tree including AIEC, commensal and pathogens strains was constructed with gff3 files provided by Prokka and used as input in Roary (Page A-L et al., 1999) 80% identity for blastp and 99% of isolates a gene must be in to be core. The tree was rooted with *Fergusonii* strains. Visualisation and customisation were performed with iTOL v6 (Letunic I & Bork P, 2024).

### Infection, viable count and antibiotic challenge

THP1 (ATCC TIB-202) monocytes (4.75x10^5^ cells/mL) differentiated into macrophages in phorbol 12-myristate 13-acetate (PMA, 20 ng/ml; Sigma-Aldrich, P1585) were infected with culture of bacteria at an OD600nm of 0.5 at MOI 100. Antibiotic challenge and viable bacterial count was performed as described before (Demarre G et al., 2019).

### Immunolabelling and Microscopy analysis

Infected THP-1 macrophages were fixed and immunostained by Lamp-I antibody (DSHB, H4A3) and WGA (Thermofisher, W32466/W32464, or Biotium, 29027-1) was performed as described previously in Prudent *et al*. Microscopy was performed as in (Demarre G et al., 2017) on an inverted Zeiss Axio Imager with a spinning disk CSU W1 (Yokogawa).

### Live-Cell Imaging

All live-cell analyses were performed using Nunc Lab-Tek II Chambered Coverglass (ThermoFisher, 155382). The acquisition was performed on an Axio Observer 7 Definite focus 2 video-microscope (Carl Zeiss Micro-Imaging) equipped with a thermostatic chamber (37°C, 5% CO2). Acquisitions were performed every 15 minutes during 16 hours. Analysis was done using FIJI software.

### FRAP analysis

FRAP was performed on THP-1 macrophages infected with LF82-GFP, CEA601S-GFP or CEA106S-GFP. Infections were conducted in fluorodish (World Precision Instruments). FRAP was undertaken at 48h P.I. Acquisitions were taken every 10 seconds using a Plan-APO 60×/1.4NA objective on a Ti Nikon microscope enclosed in a thermostatic chamber (Life Imaging Service) equipped with a Evolve EMCCD camera coupled to a Yokogawa CSU-X1 spinning disk. Four images were taken before photobleaching and fluorescence recovery was followed every 10 seconds for 5 minutes. Fluorescence recovery was collected with Metamorph Software (Universal Imaging) and was analysed using the FIJI Software.

### STED imaging

THP-1 macrophages infected by the different GFP strains were fixed and immunostained as described in Prudent *et al*. Imaging was performed with STED microscopy using an Axio Observer 7 (Carl Zeiss Micro-Imaging) equipped with a STEDYCON module (Abberior) using a x100 oil-immersion objective. Detection was realised with avalanche photodiodes with filter cubes for green (500–550 nm), red (605–625 nm), and far red (650–720 nm) fluorescence.

### Cell Viability

THP-1 macrophage viability was determined by colorimetric method using the CellTiter 96® AQueous One Solution Cell Proliferation Assay (Promega; G35C) as per the manufacturer’s instructions. Briefly, 1x105 cells were loaded into each well in a 96-well plate and infected as described before. After addition of 20µL of reagent plates were incubated and measured using TECAN Spark microplate reader. All values were normalised and compared to standard curves.

### Western Blot analysis

Cell extracts were obtained using homemade Laemmli buffer devoid of reducing reagents (200 mmol/L Tris pH 6.8, 8% SDS, 40% glycerol, 0.2% bromophenol blue) SDS-PAGE analyses were performed with 4%–12% NuPAGE bis-tris gels (Thermo Fisher Scientific, NP0336) and immunoblotted on nitrocellulose membranes (BioTrace Pall Laboratory, 732–3031), with antibodies: anti-LC3 (1/500; Sigma-Aldrich L8918), anti-Actin (1/1000; Sigma-Aldrich, A2066), anti- NFkB (1/1000; Thermofisher scientific, 51-0500), anti-RpoB (1/1000; Biolegend, 663907). Secondary antibodies provided by Jackson Immuno-Research anti-Rabbit-HRP (1/5000; 711- 035-152). Proteins were detected using SuperSignal West Pico or Femto (Thermo Fisher Scientific, 34580 or 34095), using Vilber Fusion-Fx (Vilber). Quantification was performed with FIJI software.

### FACS

Infected THP-1 macrophages were wash twice with 1X PBS, and lysed during 5 minutes with 1% Triton X-100, filtered (40µm; VWR, 732-2757) and fixed in formaldehyde (3.7% final) during 30 minutes. Samples were centrifuged, washed once and resuspended in PBS 1X. The flow cytometry measurements were obtained with CytoFlex LX cytometer and were managed using CytExpert. Analysis was performed using FlowJo Software.

### Recovery growth and CFU on supernatant

RAW 264.7 macrophages were seeded at 4,75x10^5^ macrophages/mL, 2mL/well and infected with AIEC culture at OD600 = 0.5 as described before (Prudent V et al., 2021); After 48h of infection, the macrophage supernatant is removed, and 1 mL of antibiotic-free DMEM medium is added after 2 PBS washes. 2h later, the well supernatant is collected and centrifuged; 1). For CFU count, 500µl of this supernatant is removed before resuspended cells. Next, conventional CFU counts are performed with an Easy-spiral-Interscience machine (to enumerate live bacteria in the supernatant 2). For growth evaluation, 500µl of this supernatant is removed before addition of 500µl LB medium. Growth was measured during 16 hours with Tecan Spark. Macrophages remaining in the wells were lysed with 1% Triton and CFU was performed as previously described (Prudent V et al., 2021) and counted with Scan300- Interscience software.

### Cytokines quantification

Infection in THP-1 macrophages was performed as described before. Quantification of cytokines in macrophages’ supernatant was measured using MACSPlex Cytokine kit as per the manufacturer’s protocol (Milteny, calibration: 130-106-197; TNF-α: 130-109-694; IL-6: 130- 109-691; INF-γ : 130-109-695). Flow cytometry measurements were obtained with CytoFlex

LX cytometer and were managed using CytExpert. Analysis was performed using Flowreada.io Software.

### RNA extraction and qPCR analysis

**Extraction RNA bacteria:** Total RNA bacteria extraction was performed on overnight bacteria culture; OD was adjusted to 0.2. RNA extraction was performed with Aquaphenol protocol.

**Extraction RNA of infected THP-1 macrophages:** Total RNA extraction in infected THP-1 macrophages was performed using TRIzol (Invitrogen, 15596018) and chloroform protocol. Reverse transcriptase was carried out with iScript Reverse Transcription (Biorad, 1708891) after DNase treatment (Sigma, AMPD1-1KT DNaseI) and qPCR was conducted with iQ SYBR Green Supermix (Biorad, 1708880) on 1µL of cDNA and 5µM of primers. The data was acquired on Biorad CFX Connect Real-Time System in the Biorad CFX Maestro Software.

## Acknowledgments

We are grateful to the teams of Nicolas Barnich and Margarita Martinez Medina for the strains and their precious scientific advice. We thank Eric Lallemand for his help with the project, as well as Marie-Agnès Petit, François Lecointe, and Pauline Misson for their in-depth knowledge and advice with prophages. Grateful acknowledgement also goes to Magali Fradet for her invaluable help with flow cytometry, and all the CIRB imaging platform. We would also like to thank the members of the laboratory for their careful proof-reading and suggestions on the document.

---

**Supplementary figure 1.**
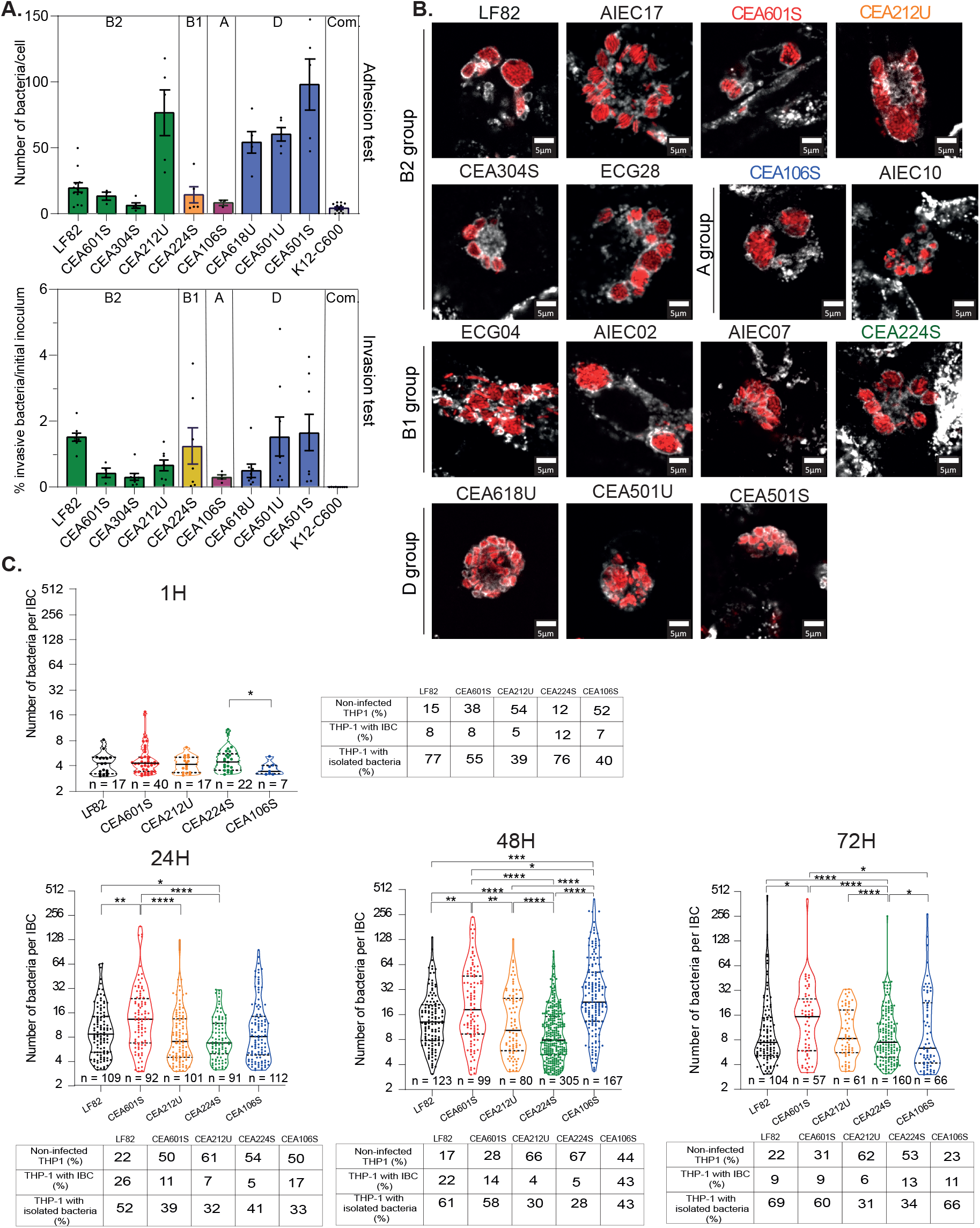
A) Adhesion assay of AIEC on epithelial cells (top) and invasion assay of AIEC into epithelial cells (bottom). B) Imaging of IBC within Lamp-I positive vacuole in THP-1 macrophages by AIECs. The scale bars are 5 µm C) For each time point in the kinetic study, a summary sheet shows the percentage of THP-1 macrophages that are non-infected, infected with IBCs, or harboring isolated bacteria. Additionally, a violin plot illustrates the number of bacteria per IBC. Data were analyzed using the Mann-Whitney test, with significance indicated as follows: *P < 0.05, **P < 0.01, ***P < 0.001, ****P < 0.0001.

**Supplementary figure 2.**
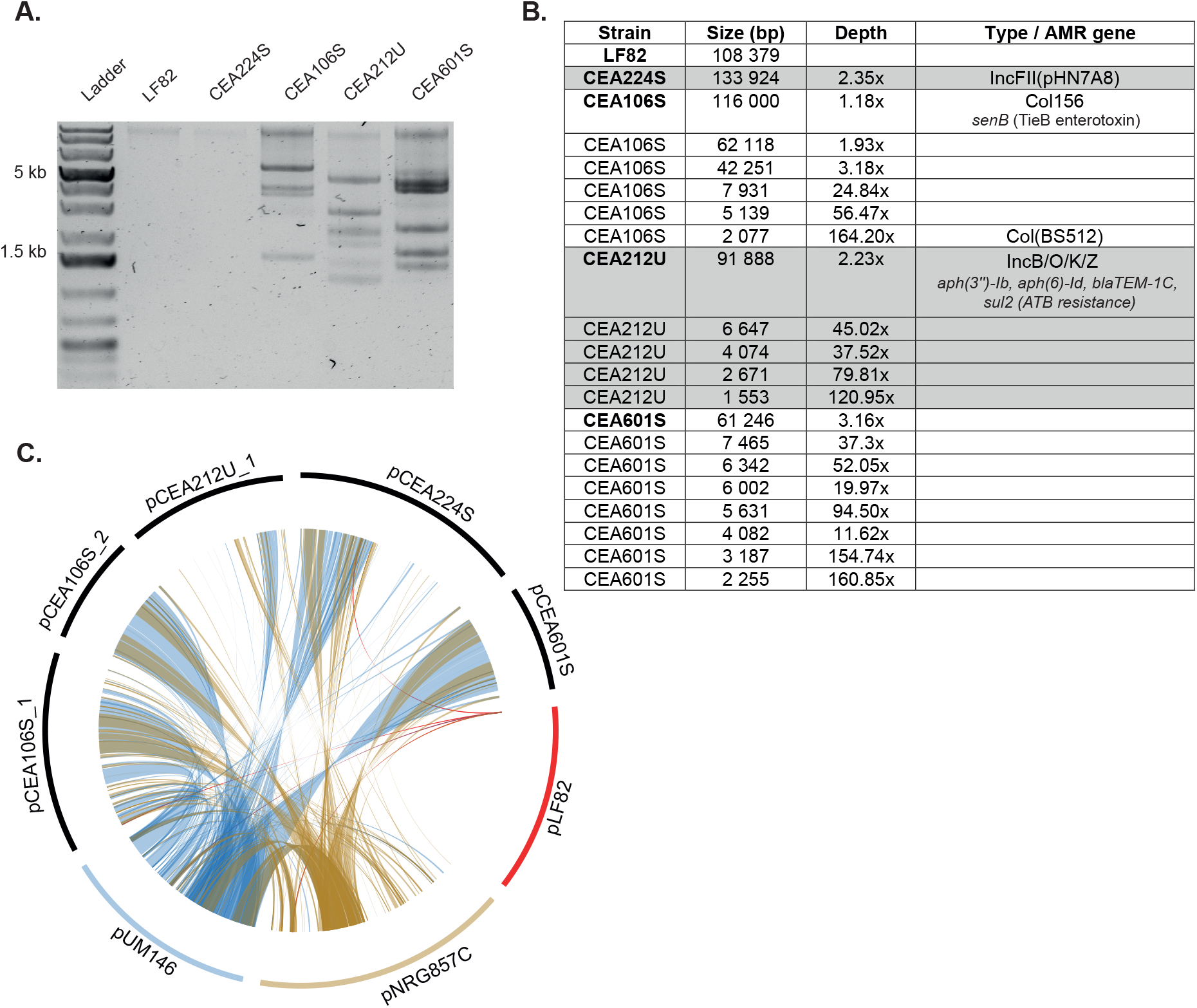
A) Plasmids of AIEC strains visualized on an agarose gel. B) Table summarizing the different AIEC plasmids, including their sizes, sequencing depths, and types. The coding sequences (CDS) of plasmids pLF82, CEA106Spa.1, CEA106Spb.1, CEA106Spc.1, CEA212Up.1, CEA224Sp.1, and CEA601Sp.1 are detailed in Table S2. C) Circos plot comparing AIEC plasmids from the five studied AIEC strains, as well as pUM146 and pNRG857C.

**Supplementary figure 3.**
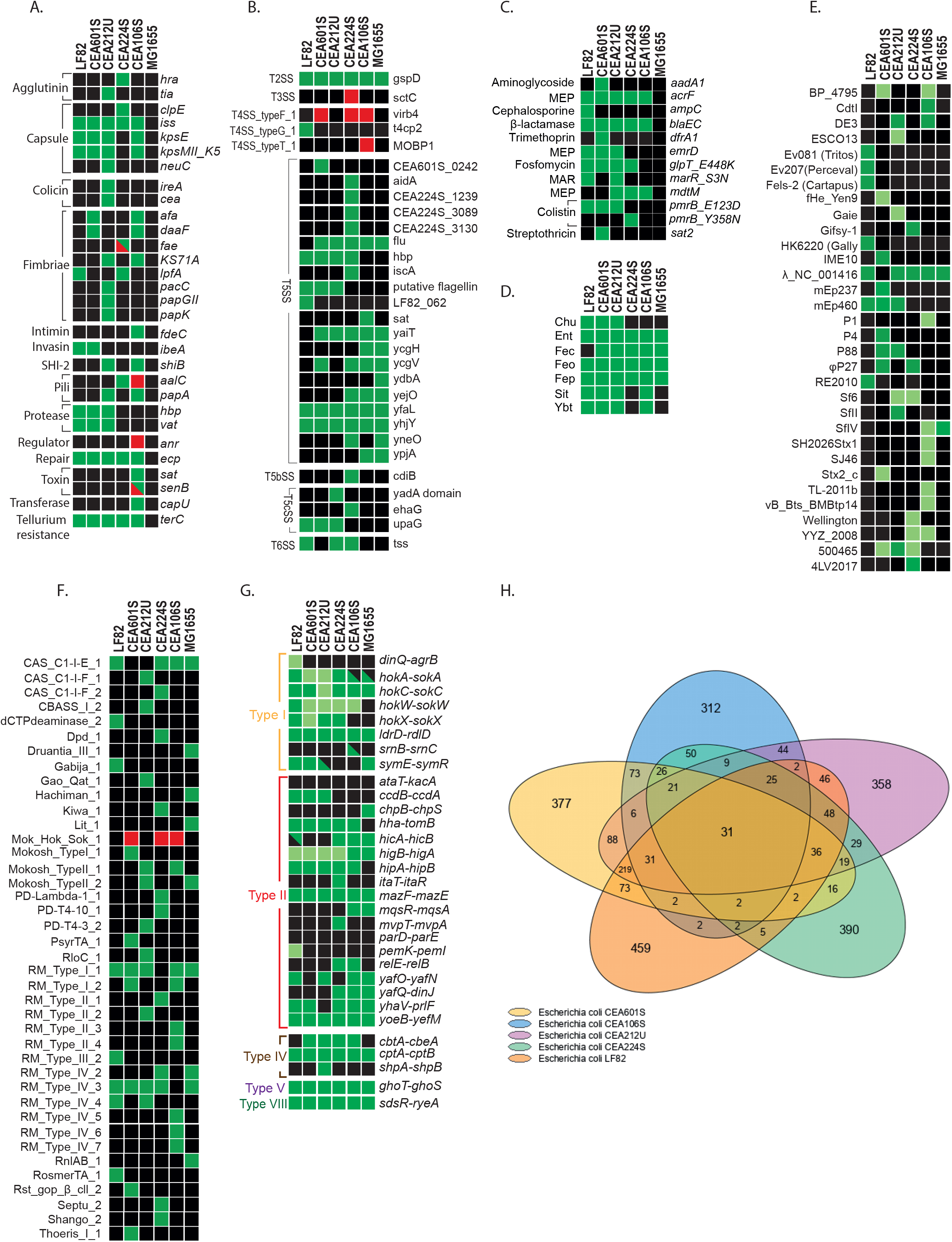
Presence/absence matrix of genes for the five AIEC strains and MG1655 K12. The color coding is consistent across all graphs: green squares indicate the presence of a gene in the chromosome, while black squares denote its absence. Red squares signify the presence of the gene on a plasmid. For systems with multiple genes, light green squares represent partial presence. A) Matrix of predicted virulence factors based on VFDB. B) Matrix of predicted secretion systems identified using MacSyFinder on the Mage Genoscope website. C) Matrix of predicted antimicrobial resistance based on AMRfinder. D) Matrix of iron acquisition systems. E) Matrix of predicted prophages using Phigaroin on Mage. F) Matrix of defense system predicted by CRISPRCasFinder in Mage Genoscope website. G) Matrix of toxin-antitoxin system predicted by TAfinder2.0. H) Venn Diagram of the core genome of the 5 AIECs without the gene present in MG1655 K12 genome.

**Supplementary figure 4.**
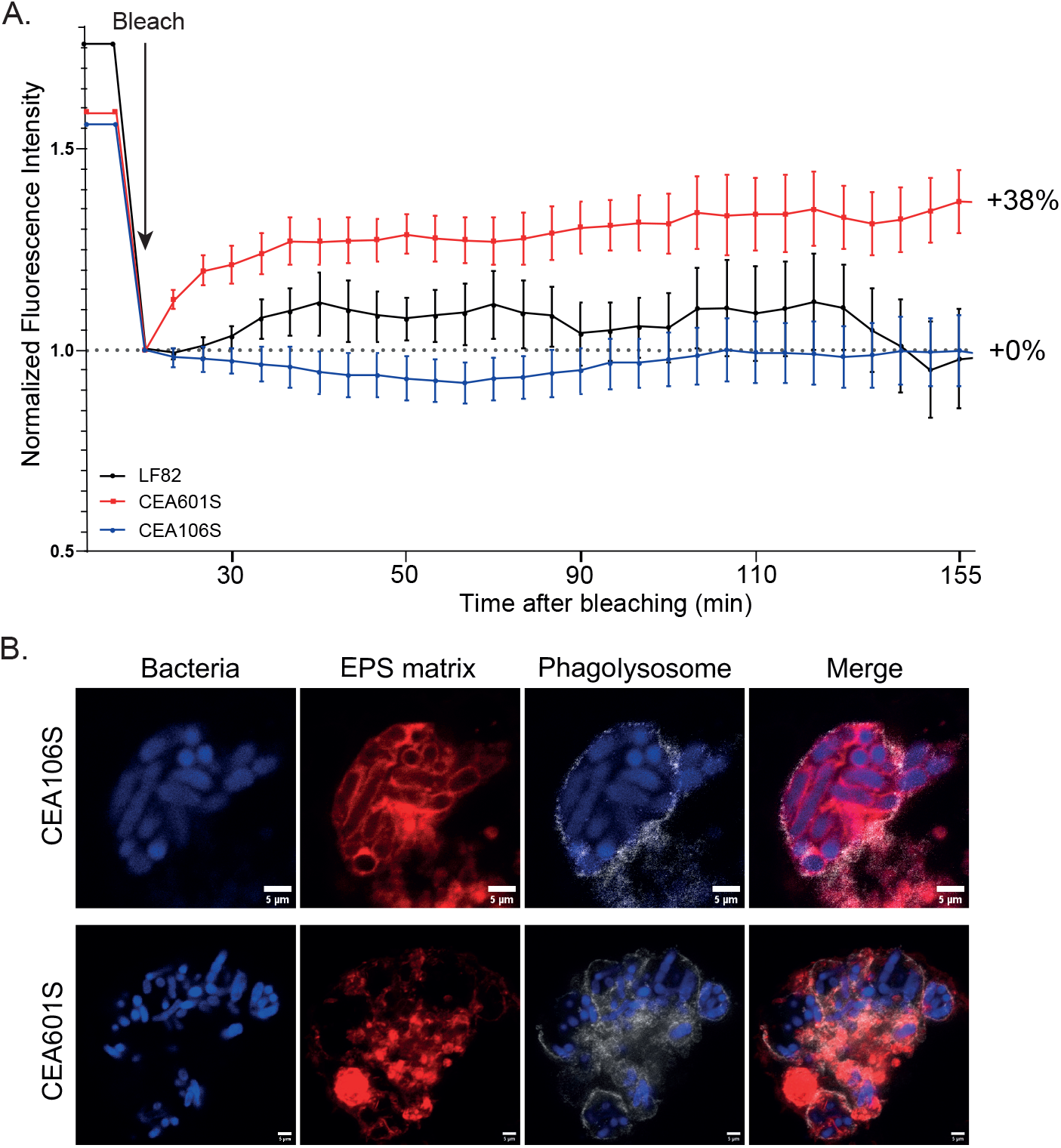
A) FRAP experiments on phagolysosomes containing LF82-GFP, CEA601S-GFP, or CEA106S-GFP, conducted at least 48 hours post-infection (P.I.). Data after bleaching were normalized to 1 to assess fluorescence recovery. B) STED imaging of IBCs within THP-1 macrophages at 24 hours P.I. Bacteria are shown in blue, exopolysaccharide (EPS) matrix is labeled with WGA lectin in red, and phagolysosomes are labeled with Lamp-I antibody in grey. Scale bars represent 5 µm.

**Supplementary figure 5.**
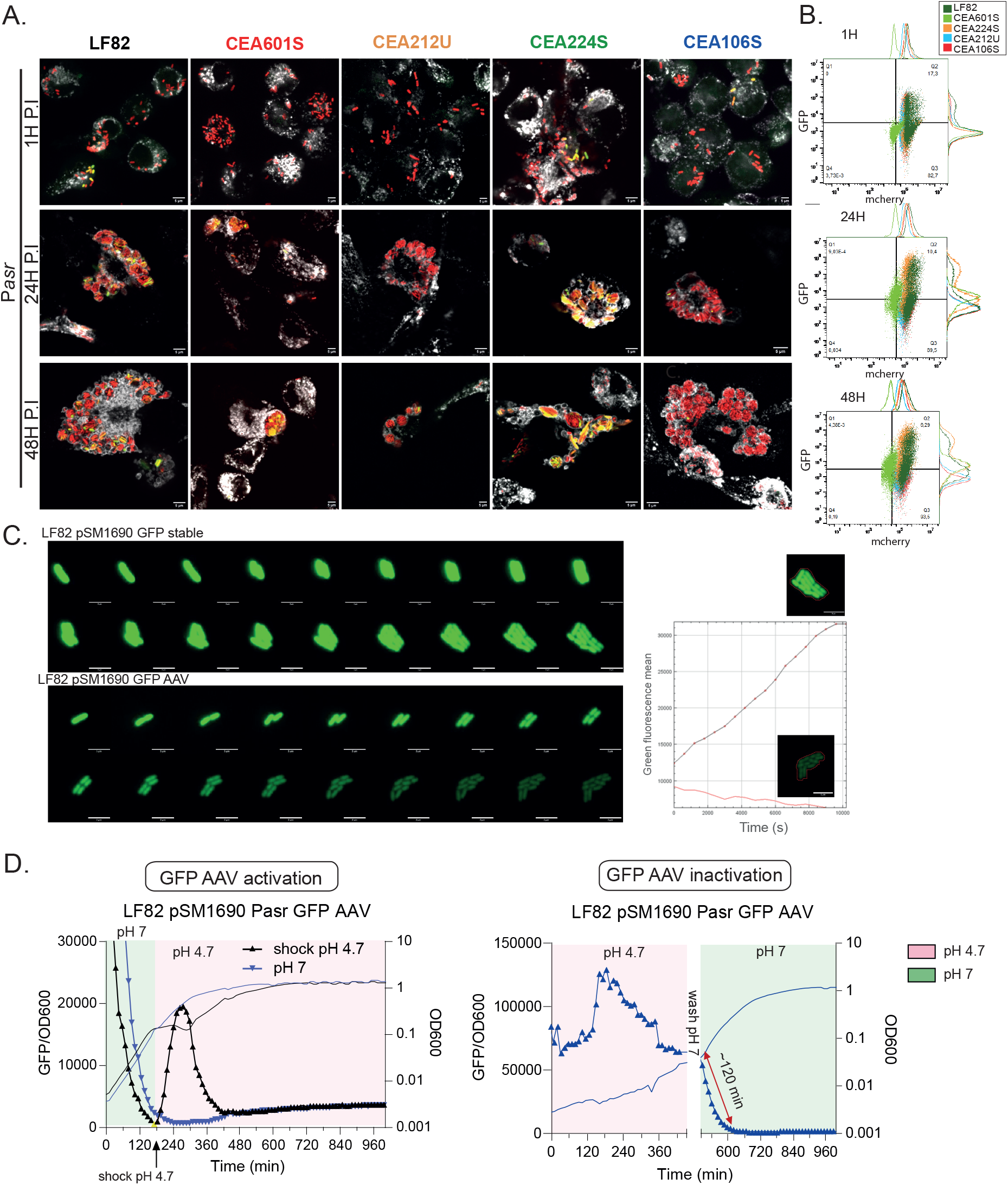
A) Imaging of THP-1 macrophages infected with AIEC strains carrying a plasmid reporter for asr (acid response) promoter activation, with unstable GFP. Imaging was performed at 1, 24, and 48 hours post-infection (P.I.). Bacteria are shown in red, with green indicating activation of Pasr. Phagolysosomes are labeled with Lamp-I antibody and appear in grey. Scale bars represent 5 µm. B) Histogram from FACS analysis of the asr reporter at 1, 24, and 48 hours P.I. C) *In vitro* validation of unstable GFP compared to stable GFP for the asr reporter over time, including microscopy images and quantification of GFP intensity. Scale bars are 5 µm. D) *In vitro* validation of unstable GFP activation (left) and inactivation (right) with fluorescence monitoring using a microplate reader.

**Supplementary figure 6.**
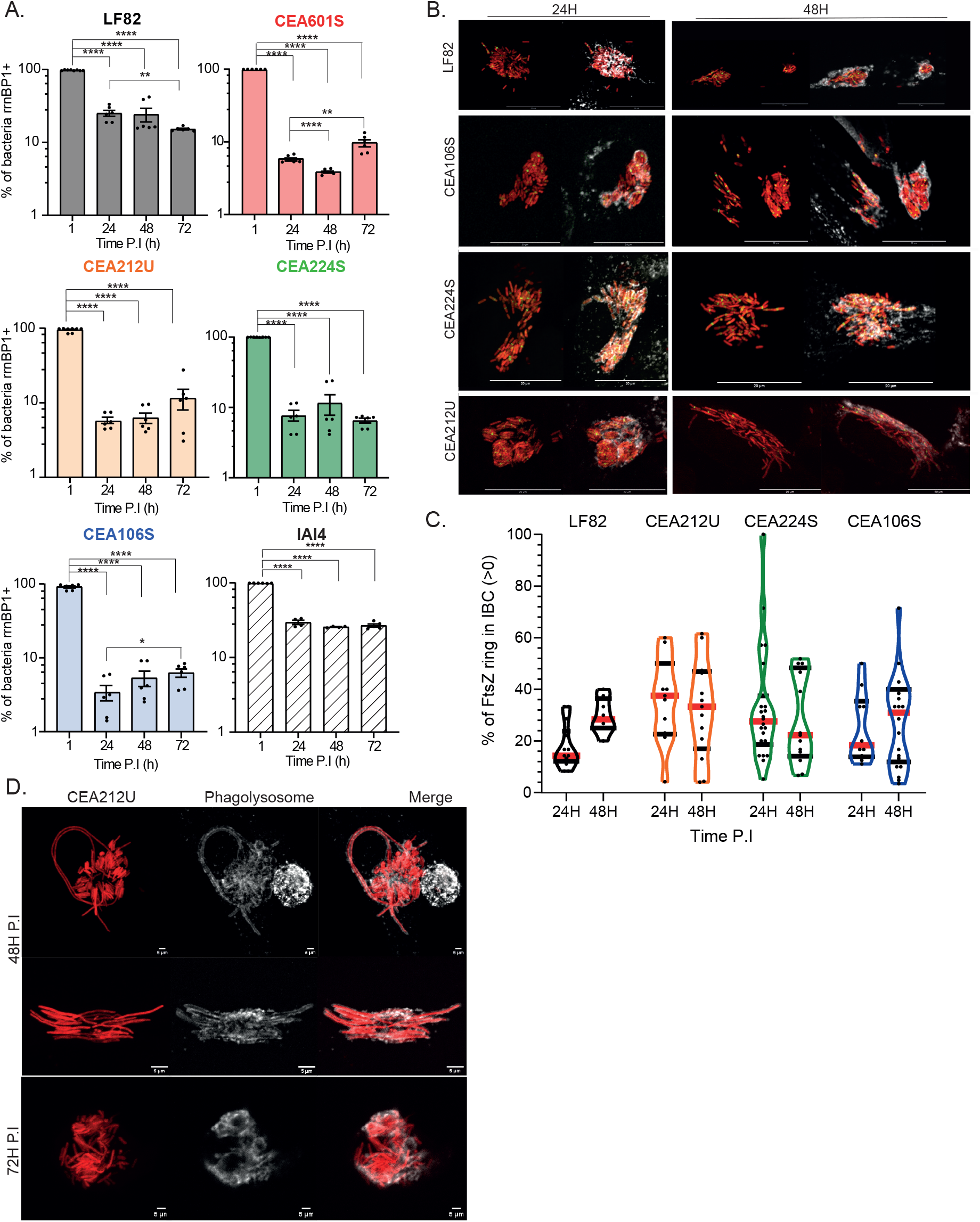
A) Imaging of filamentous CEA212U bacteria within THP-1 macrophages at 48 and 72 hours post-infection (P.I.). Bacteria are shown in red, and phagolysosomes are labeled in grey using Lamp-I. Scale bars represent 5 µm. B) Imaging of AIEC strains containing an FtsZ-GFP reporter within Lamp-1 vacuoles (grey) in THP-1 macrophages at 24 hours P.I. (left panel) and 48 hours P.I. (right panel). Scale bars represent 20 µm. C) Quantification of FtsZ-positive bacteria within IBCs at 24 and 48 hours P.I. D) Flow cytometry analysis of AIEC response to intracellular nutrient scarcity (*rrnBP1*). P-values are indicated (Student’s t-test: *P < 0.05, **P < 0.01, ***P < 0.001, ****P < 0.0001); error bars represent the mean and standard error of the mean (SEM); n ≥ 3. Flow cytometry gating strategies are described in Supplementary Figure 8.

**Supplementary figure 7.**
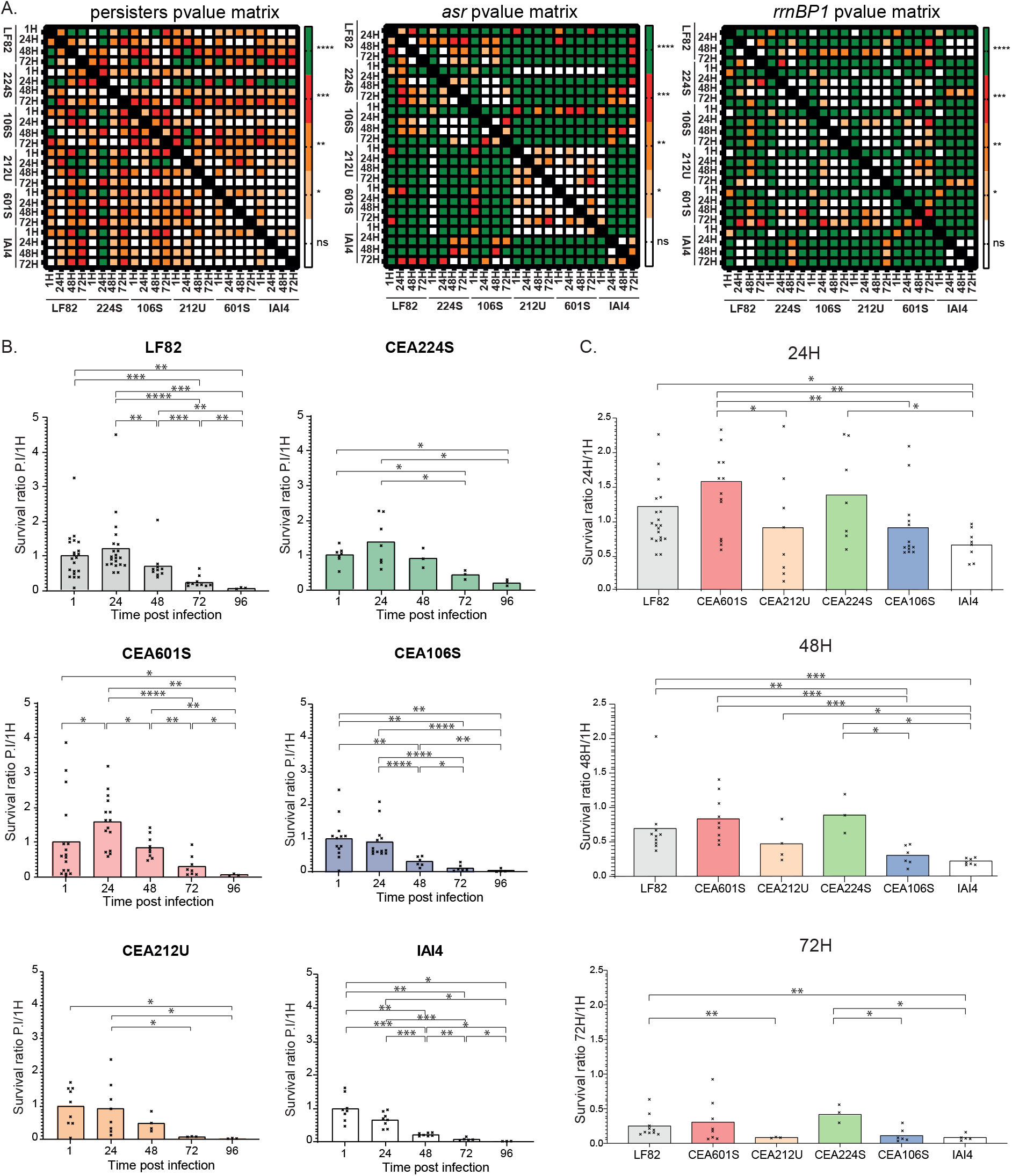
A) Matrix of p-values for persisters (left panel), *asr* FACS analysis (center panel), and *rrnBP1* FACS analysis for strain comparison. Data were analyzed using Student’s t-test: *P < 0.05, **P < 0.01, ***P < 0.001, ****P < 0.0001. B) Proportion of viable bacteria during the infection kinetics in THP-1 macrophages compared to levels at 1 hour post-infection (P.I.). Each graph represents a single strain. P-values are indicated (Mann-Whitney test: *P < 0.05, **P < 0.01, ***P < 0.001, ****P < 0.0001). C) Proportion of viable bacteria in THP-1 macrophages at different time points compared to 1 hour P.I. Each graph represents a different time point. P- values are indicated (Mann-Whitney test: *P < 0.05, **P < 0.01, ***P < 0.001, ****P < 0.0001).

**Supplementary figure 8.**
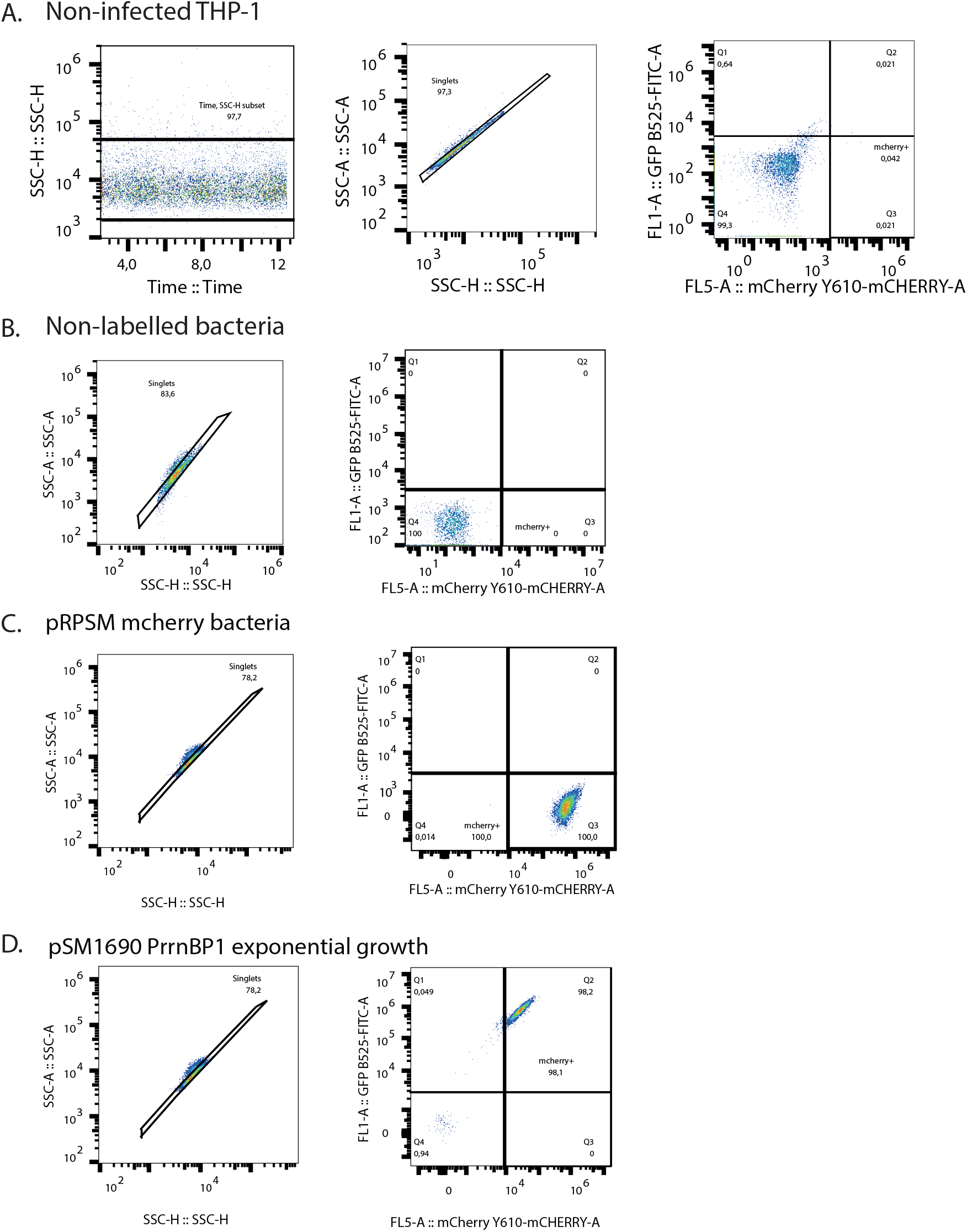
Flow cytometry gating strategies for quantifying populations activating the asr and rrnBP1 promoters. A) Gating of non-infected and lysed THP-1 macrophages. B) Gating on exponentially growing unlabelled bacteria to determine fluorescence thresholds. C) Gating on exponentially growing bacteria with a constitutive red plasmid to establish the red fluorescence axis. D) Gating on exponentially growing bacteria with the rrnBP1 plasmid to establish the green fluorescence axis. For all experiments, the Q3 quadrant (bottom right) and Q2 quadrant (top right), which correspond to red-positive events, are combined to obtain the total bacterial population. The number of events in the Q2 quadrant (top left), corresponding to green-positive events (indicative of promoter activation), is then divided by the total red-positive events.

